# Deep-LUMEN Assay – Human lung epithelial spheroid classification from brightfield images using deep learning

**DOI:** 10.1101/2020.10.07.328005

**Authors:** Lyan Abdul, Shravanthi Rajasekar, Dawn S.Y. Lin, Sibi Venkatasubramania Raja, Alexander Sotra, Yuhang Feng, Amy Liu, Boyang Zhang

## Abstract

Three-dimensional (3D) tissue models such as epithelial spheroids or organoids have become popular for pre-clinical drug studies. However, different from 2D monolayer culture, the characterization of 3D tissue models from non-invasive brightfield images is a significant challenge. To address this issue, here we report a Deep-Learning Uncovered Measurement of Epithelial Networks (Deep-LUMEN) assay. Deep-LUMEN is an object detection algorithm that has been fine-tuned to automatically uncover subtle differences in epithelial spheroid morphology from brightfield images. This algorithm can track changes in the luminal structure of tissue spheroids and distinguish between polarized and non-polarized lung epithelial spheroids. The Deep-LUMEN assay was validated by screening for changes in spheroid epithelial architecture in response to different extracellular matrices and drug treatments. Specifically, we found the dose-dependent toxicity of Cyclosporin can be underestimated if the effect of the drug on tissue morphology is not considered. Hence, Deep-LUMEN could be used to assess drug effects and capture morphological changes in 3D spheroid models in a non-invasive manner.

**Significance of the work:** Deep learning has been applied for the first time to autonomously detect subtle morphological changes in 3D multi-cellular spheroids, such as spheroid polarity, from brightfield images in a label-free manner. The technique has been validated by detecting changes in spheroid morphology in response to changes in extracellular matrices and drug treatments.

## Introduction

Advances in biological research rely on the use of effective *in vitro* culture systems. By far, the most commonly used approach is to culture primary cells or cell lines on 2D surfaces in a multi-well plate. However, it is generally recognized that the flat and hard plastic substrates commonly used are not representative of the cellular environment found in organisms. For instance, studies have shown that epithelial cells on monolayer culture have compromised integrin function leading to a higher frequency of chromosome mis-segregation during proliferation^1^. On the contrary, growing cells within a natural extracellular matrix that permits self-organization in 3D can significantly improve chromosome segregation fidelity^1^. Both primary chondrocytes and hepatocytes have also been shown to lose their normal phenotype rapidly once removed from the body and when placed in 2D culture^2,3^. Issues like these significantly limit the potential of 2D culture systems to predict the cellular responses of real organisms.

Recognizing these challenges, 3D tissue culture with tissue-specific architecture, mechanical and biochemical cues, and cell-cell communication could help reduce the gap between cell-based assays and physiological tissues^4^. Stem-derived organoids or 3D spheroids are self-organized multi-cellular tissue grown in 3D hydrogel matrices. Within this 3D environment, cells can sense a substrate that more closely resembles native extracellular matrices and have the freedom to remodel and form 3D organ-specific structures. 3D cancer spheroids have been shown to outperform 2D cell monolayers in drug screening^5^. Various organ-specific organoids from the kidney^6^, colon^7^, brain^8^, and liver^9^ have shown sophisticated tissue functions that would be impossible to replicate in 2D. For these reasons, 3D spheroid and organoid cultures are fast becoming the ideal model systems for *in vitro* drug testing and biological research.

Despite these advantages, there are barriers to using 3D models for pre-clinical drug testing. High-throughput screening has been optimized for monolayer culture for decades. 3D tissue models often lack automated workflows for analysis with fast processing times^9^. More importantly, characterizing 3D tissue morphological changes in response to drug treatments, even though it can provide highly relevant physiological information that cannot be easily derived from 2D culture, is challenging. Conventional imaging processing techniques, mostly suitable for 2D cultures, fall short in accurately characterizing complex 3D features that are not a simple description of the area, size, or shape, which can be more easily defined. In addition, the 3D environment presents artifacts like overlapping tissues, out of focus tissues, varying light conditions, tissue heterogeneity, or even supporting tissues like vasculatures, which all present significant challenges to conventional imaging processing techniques.

Deep learning could overcome these challenges by circumventing the need to arbitrarily define multiple parameters for any given set of images as in the conventional imaging processing technique. Specifically, convolutional neural networks (CNNs) are a class of deep learning and are mainly used for image analysis. CNNs recognize patterns from large training datasets, emulating the learning process inherent to our brain; hence it does not require any parameter tuning and runs autonomously. A truly myriad of applications has been explored using this technology. For spheroid and organoid analysis, machine and deep learning have been previously used to (1) localize spheroids and organoids in 3D cultures and determine their diameter^10-12^ and (2) to segment spheroids^11^. Although epithelial spheroid polarity has been classified by deep learning, this analysis was done on fluorescent images^13^. Deep learning has yet to be applied to detect subtle changes in 3D multi-cellular tissue morphology, such as spheroid polarity, from brightfield images in a label-free manner.

Utilizing Tensorflow Object Detection API^14^, here we present an open-source algorithm, Deep-Learning Uncovered Measurement of Epithelial Networks (**Deep-LUMEN**), which has been trained by a large dataset to enable users to detect changes in the luminal structure of 3D spheroids automatically. The formation of the lumen is an indication of proper cell polarization, and its disruption is a significant phenotypic change in many tissues. In this validation study, we first trained Deep-LUMEN to classify the polarity of spheroids directly from brightfield images. Then, we validated this algorithm by tracking how spheroid morphology changes in response to different types of extracellular matrices and drug treatment.

## Results

### Lung alveolar spheroid generation

Lung alveolar epithelial cells (A549), when embedded in a 3D hydrogel matrix (such as Matrigel^®^ or Fibrin gel), can proliferate and self-organize into a 3D spheroid over time. However, influenced by the culture environment, only a fraction of the population will self-organize into spheroids with a hollow lumen, which is the correct morphology of lung alveoli. Therefore, tracking this morphological feature could allow us to predict the changes in the health and function of the lung alveolar epithelium in response to environmental cues and drug treatment (**Figure 1a**). Utilizing this self-assembly process, we developed the lung spheroid models by casting single-cell suspension in Matrigel^®^ in standard 384-well plates (**Figure 1b**). The high-throughput 384-well plate, coupled with a high-content image cytometer, allowed us to automate the image acquisition process and collect thousands of brightfield images. Images were taken at different planes (spaced 50 μm apart) through the z-axis to capture all the spheroids in the scanned volumetric space (**Figure 1c**).

**Figure 1.**
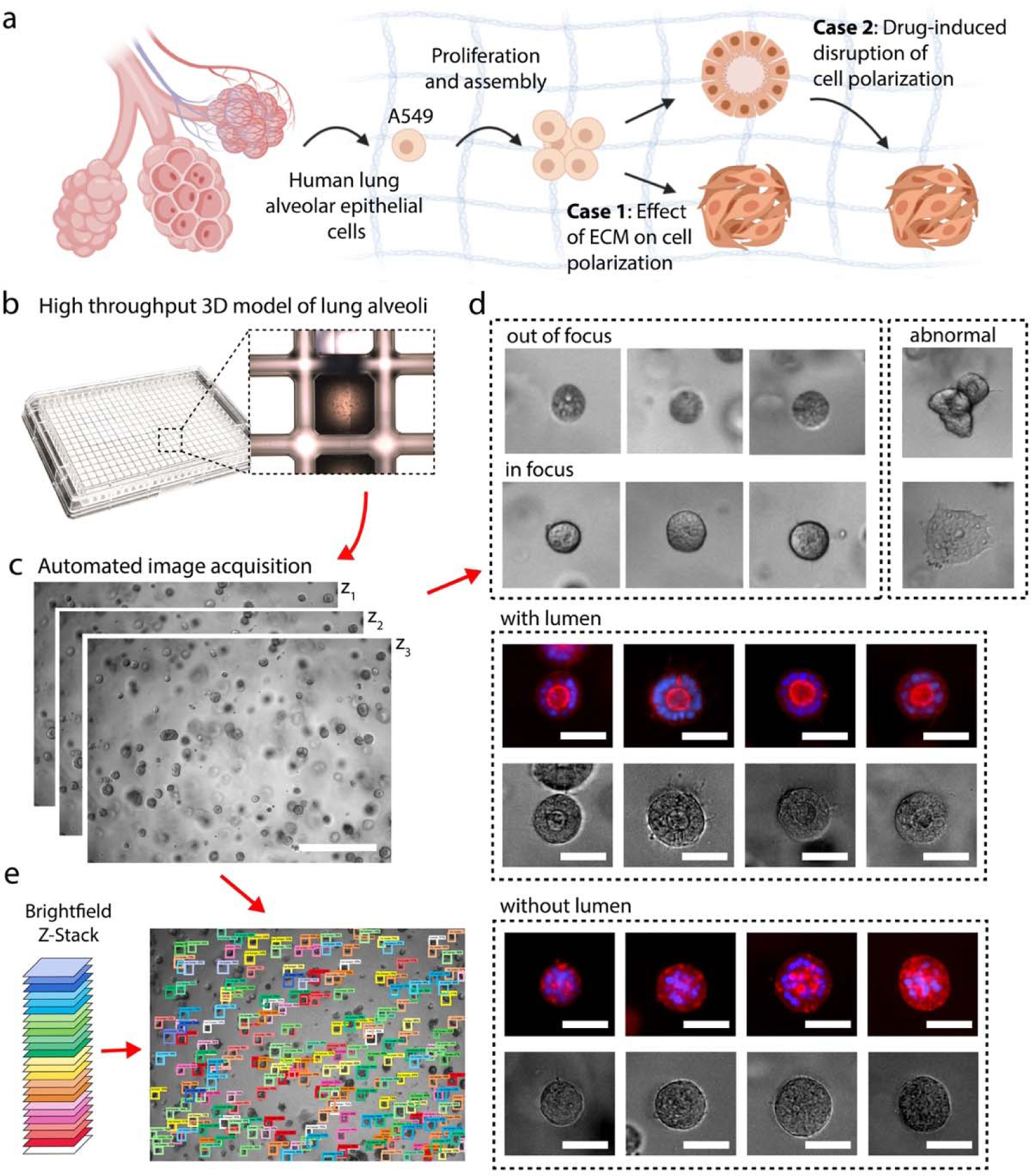
Differential formation of hollow lung alveolar spheroids. **a**, Illustration of lung epithelial cells proliferating and assembling into either hollow or solid spheroids in a 3D matrix. **b**, Tissue culture setup where 25μL of Matrigel^®^ embedded with cells are cast in standard 384-well plates. **c**, acquired z-stack transmission-light images. **d**, Example scenarios of lung spheroids seen from the collected images. Corresponding fluorescent images stained for F-actin (red) and DAPI (blue) of lung spheroids with or without a lumen (representative images from n=6 samples). Scale bar, 50 μm. **e**, z-stack acquisition allows for spheroid morphology assessment throughout entire hydrogel. Spheroids on different focal planes were detected with developed Deep-LUMEN algorithm from z-stack images and then labeled with different colors for visualization.

### Training dataset generation

We created a custom training dataset containing 4000 images (14,993 examples of no lumen and 2,351 examples of lumen spheroids). Out of focus spheroids were omitted from each image to avoid double-counting. Any abnormal morphologies or monolayer formations were also omitted (**Figure 1d**). Bounding boxes were drawn around the in-focused spheroids and categorized accordingly. A spheroid was categorized as “lumen” if there was a prominent indent in the center of the spheroid. Otherwise, the spheroid was given a “no lumen” classification. Immunofluorescent staining confirms that lumen formation is directly correlated to this indent feature from brightfield images and also correlates to cell polarization and the proper organization of the lung alveolar cells. Stronger F-actin staining was seen on the apical side of the luminal spheroid, while disorganized spheroids showed intense F-actin staining on both basal and apical sides (**Figure 1d, Supplementary Figure 1**). It’s important to note that even though the immunofluorescent staining validated our assumption, the immunofluorescent assay is terminal. On the contrary, Deep-LUMEN classification from brightfield images is non-invasive and continuous. In addition, changes in spheroid morphology can be automatically assessed for each image present in the z-stack, providing a characterization of spheroid morphology through the entire hydrogel (**Figure 1e**). The full training dataset, including the images and annotations, are publicly available at https://osf.io/g2a7r/

### Model optimization

After training, the performance of the algorithms were then tested on a set of one-hundred and ninety-seven new images. The models outputted bounding boxes around spheroids along with the classification and the confidence probability. Furthermore, the algorithms learned to omit out-of-focus spheroids and spheroids of abnormal shape (**Figure 2a**). To get the best results, we tested five different pre-trained models and fine-tuned them on our lung spheroid dataset^15-18^. Out of the five models (Model 1-5, **Figure 2b**), Faster R-CNN with ResNet101 (Model 5) performed the best. When tested with a new test-set (197 new images) it had a 73% mean average precision (**Figure 2d**) along with the fewest false positives and false negatives (**Figure 2c**). To further improve this model’s performance, we then added additional data augmentation options to generate Model 6. When tested with the test-set images, Model 6 (Faster R-CNN ResNet101 with data augmentation) had a 83% mean average precision (**Figure 2e**), detected the highest number of “lumen” and “no lumen” spheroids, and had a lower rate of false positives and false negatives (**Figure 2c**). Therefore, this Model 6 was used at the end and is referred to as Deep-LUMEN. Overall, the analysis and classification of spheroids using Deep-LUMEN were more than 20 times faster than manual labeling (**Figure 2f**). Additionally, Deep-LUMEN outputs confidence scores for each of its predictions. We observed that the score corresponded with the extent of lumen development. For the non-polarized spheroids, the confidence decreases when an indentation is present. For the polarized spheroids, the confidence increases as the indentation becomes larger and the lumen is more well-defined (**Figure 2g**). This observation indicates that it could be possible to further stratify the stages of spheroid lumen development based on the confidence score.

**Figure 2.**
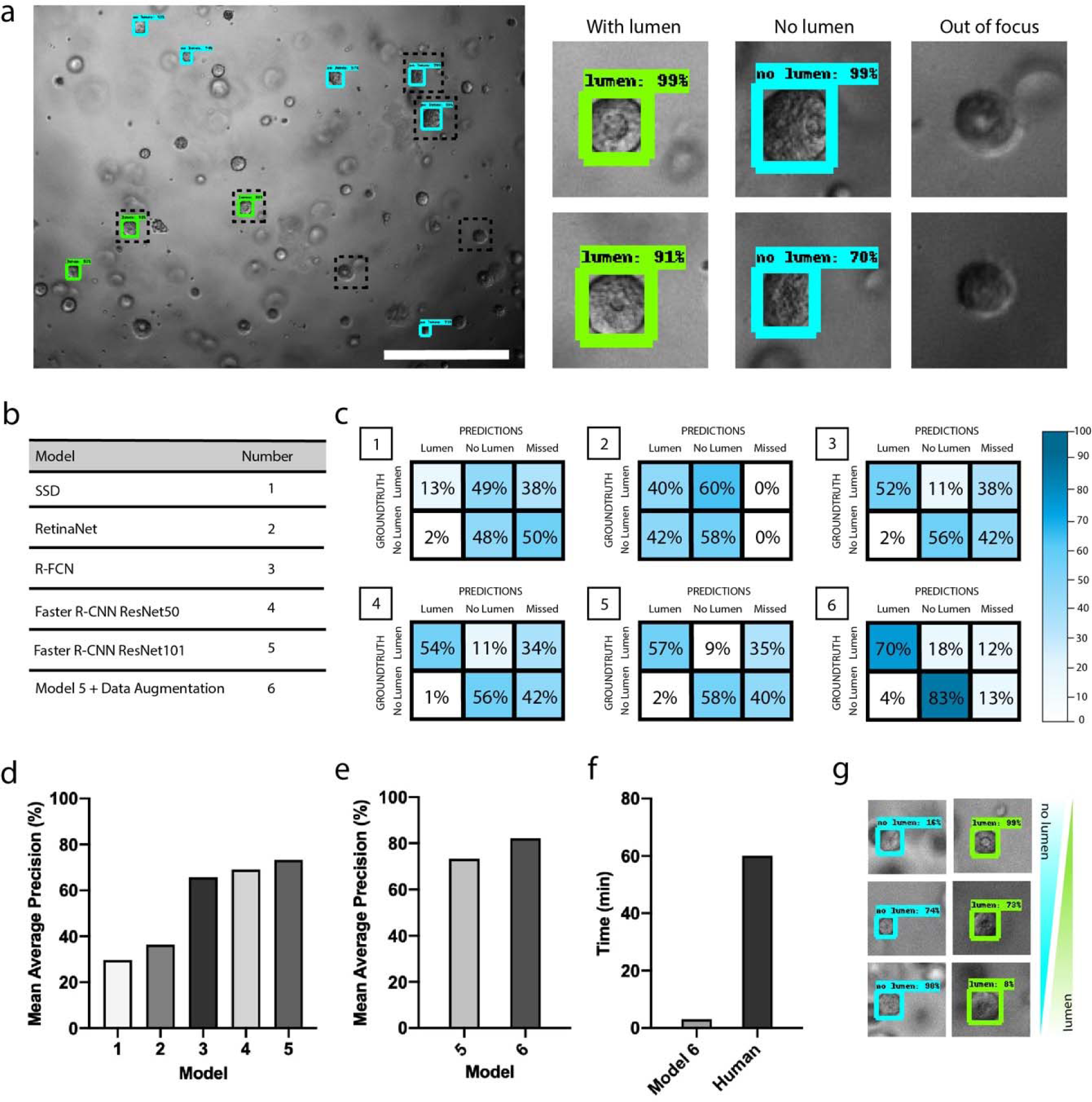
Optimization of model performance. **a**, Detection output of Deep-LUMEN following fine-tuning with custom dataset. Bright-field images were obtained by taking 4X z-stack images of lung spheroids cultured in Matrigel^®^. Scale bar, 500 μm. **b**, Models used and their corresponding Model number. **c**, Confusion matrix for each trained model outlining the true positives, the true negatives, the false positives and the false negatives. Assessment of accuracy was performed on 197 new test images. **d**, mAP metric for each fine-tuned model. **e**, Comparison of model 5 performance with and without data augmentation. Model 6 (model 5 with data augmentation) was chosen as the final model and will be referred to as Deep-LUMEN. **e**, Comparison of the time required to localize and classify spheroids in 25 images between Deep-LUMEN (on a GPU) and human annotators. **g**, Deep-LUMEN’s confidence scores reflect the extent of lumen formation.

### Spheroid morphological changes in response to extracellular matrices

We then used Deep-LUMEN to assess the morphological changes of lung spheroids in response to different extracellular matrices (ECM). Lung alveolar epithelial spheroids were embedded in four different hydrogel conditions and cultured for 12 days in a standard 384-well plate (**Figure 3a**). In all conditions, the alveolar epithelial cells started to proliferate, and the diameter of the spheroids increased over time (**Figure 3b)**. There were no significant differences between the different groups. However, the percentage of polarized spheroids (spheroids containing a lumen) in 10 mg/mL fibrin on Day 8 was significantly lower than in the Matrigel^®^ condition (**Figure 3c**). This suggests an interplay of signals between the Matrigel^®^ and the epithelial cells allowing for the formation of the lumen by guiding epithelial polarization. It is well known that laminin, a major component of the basement membrane and also abundant in Matrigel^®^ are integral for guiding cells to develop the polarity required for lumen formation^19^. Therefore, we found Matrigel^®^ to be optimal for lung spheroid culture. The unique aspect of this study is that we were able to make this conclusion by analyzing just the brightfield images with Deep-LUMEN in a completely non-invasive manner. It is also clear that we would not be able to capture this effect should we have solely focused on quantifying spheroid diameters using conventional image analysis methods.

**Figure 3.**
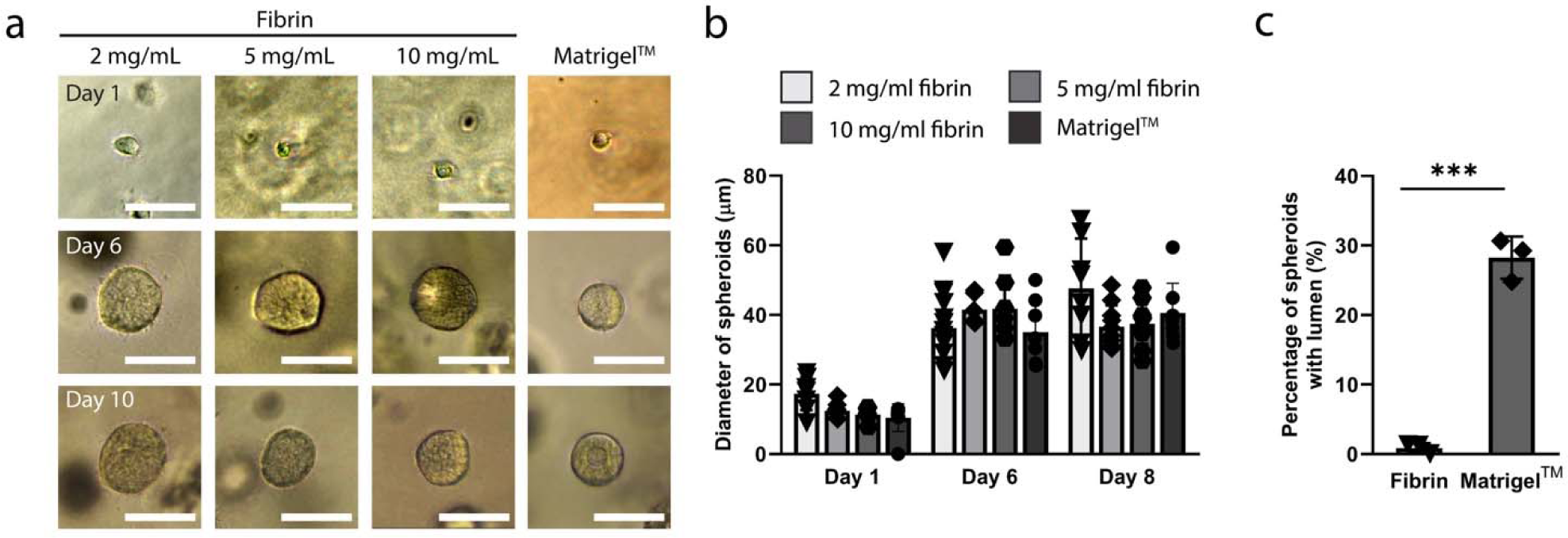
Spheroid morphological changes in response to extracellular matrices. **a**, Assembly of lung alveolar epithelial cells into spheroids in different ECM conditions. **b**, Quantification of the diameter of spheroids in different ECM conditions (n= 6 to 18 spheroids). No significant differences were found. Statistical significance was determined using one-way ANOVA on ranks with Dunn’s method. **c**, Quantification of the percentage of spheroids with the presence of lumen using Deep-LUMEN in 10 mg/ml fibrin and Matrigel^®^ (n=3). Statistical significance was determined using one-way ANOVA with the Holm-Sidak method. *p < 0.05, **p < 0.01, ***p < 0.001.

### Spheroid morphological changes in response to drug treatment

Lastly, we examined how drug treatment could affect lumen formation in the lung alveolar spheroids. Cyclosporin A (CsA) is an immunosuppressant medication used after organ transplantation^20^ and for other autoimmune conditions such as rheumatoid arthritis^21^, psoriasis^22^, etc. However, at higher dosages, it is known to be toxic to kidney or liver cells. Many studies have shown that a dosage above 10 μM (around 100 μM) is required to observe a significant toxic effect from CsA on liver or kidney epithelial cells in vitro^23,24^. Here we tested CsA over a wide concentration range from 0.01 to 10 μM (**Figure 4a**). We found that at 10μM the ability of the cells to self-organize into a hollow alveolar spheroid is already compromised, even though there is no significant effect on the size and number of lung spheroids due to CsA which indicates the drug is yet to compromise cell viability (**Figure 4 b-d**). While spheroids will, by default, develop more lumens over time, the group treated with 10μM CsA failed to form more luminal spheroids and resulted in a drastic decrease in luminal spheroid count compared to the control three days after drug administration (**Figure 4e**). This suggests that the effective toxicity of CsA could be much lower than initially expected. This further highlights the need to examine higher-level tissue morphological changes in addition to the more obvious toxicity effect on cell viability and growth in drug testing.

**Figure 4.**
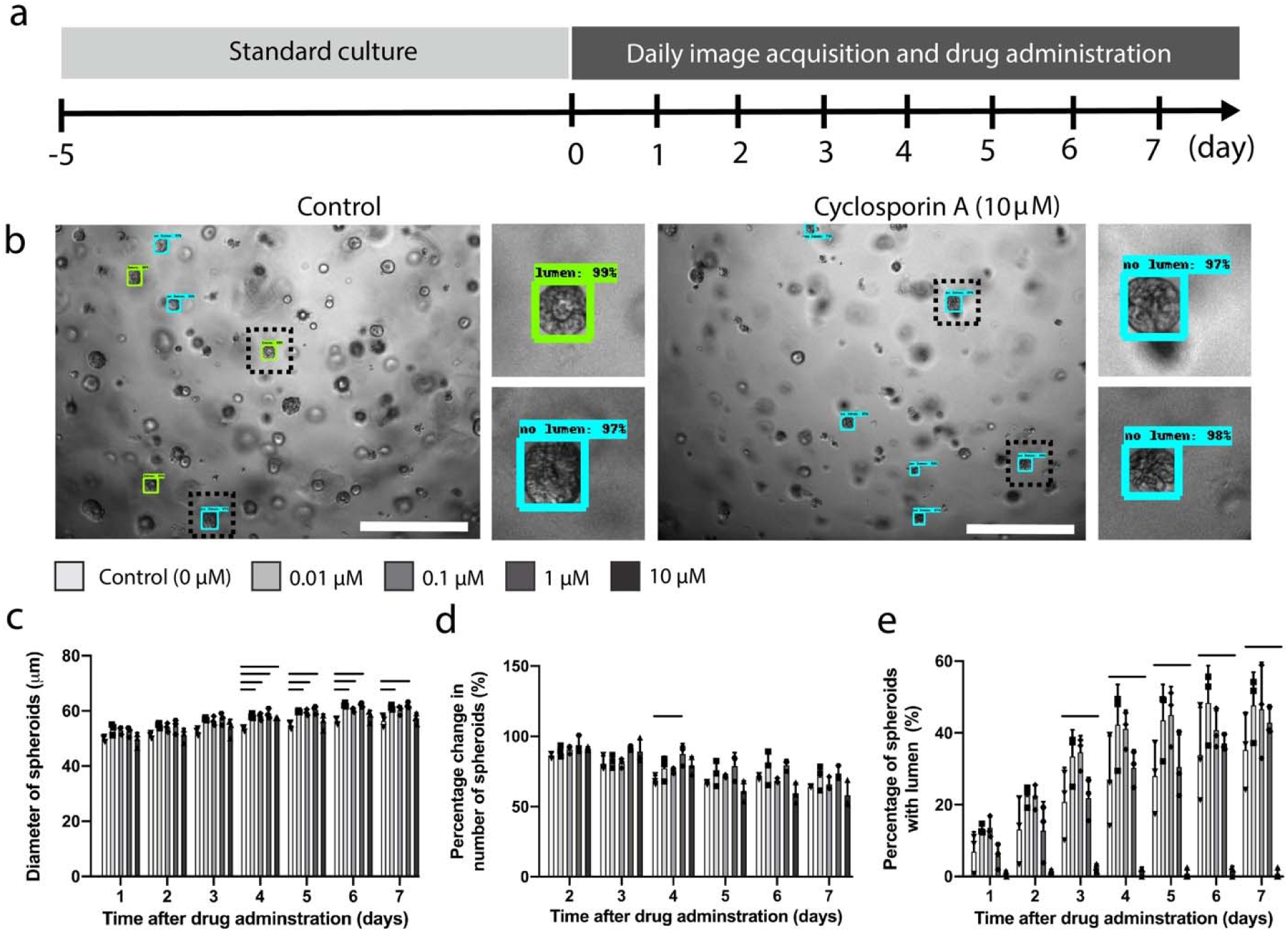
Validation of Deep-LUMEN with drug administration. **a**, Cyclosporin A administration timeline. Cyclosporin A was administered on the sixth day of lung spheroid culture for a period of seven days. **b**, Comparison of detections in control versus 10 μM drug-treated conditions. Deep-LUMEN was used to detect polarized and non-polarized spheroids following drug administration. Bright-field images were acquired by taking z-stack images of spheroids under 4X magnification (n = 3). Scale bar, 500 μm. **c**, Average diameter of spheroids following drug administration (n = 3). Bounding box coordinates of detections above a 50% score confidence were outputted to measure the diameter. Statistical significance was determined through a one-way ANOVA test followed by the Holm-Sidak method. Line indicates a p-value < 0.05. **d**, Percent change in the number of spheroids following a period of drug administration. Quantification was done by outputting detections that had at least a 50% score confidence. Statistical significance was determined through a two-way ANOVA test followed by the Holm-Sidak method. Line indicates a p-value < 0.05. **e**, Proportion of the lumen class of spheroids over the period of drug administration (n = 3). Quantification of lumen spheroids was obtained by filtering detections with at least a 50% score confidence that were of the lumen class. Statistical significance was determined through a one-way ANOVA test followed by the Holm-Sidak method. Line indicates a p-value < 0.05.

## Discussions

Object detection and classification, primarily based on subtle changes in morphological features, is a challenging task but needed if we want to fully explore the benefits of 3D models in biological research. In this study, we hope to lay the groundwork to help further apply machine learning to analyze more complex tissue models. Specifically, we showed that by offering a closer look at the changes in tissue morphology, deep learning could help improve the sensitivity of tissue models to drug treatments.

Because the Deep-LUMEN algorithm was not just localizing the spheroids, it’s important to note that the collection of z-stacked brightfield images and the ability to eliminate out-of-focus spheroids is critical for avoiding double counting and for accurately classifying the spheroids, as out-of-focus spheroids will not contain sufficient morphological details to allow accurate characterization. The z-stack scanning capability is a common feature in most high-content cytometers used for high-throughput screening. Also, the Deep-LUMEN program does not require any preprocessing on the images, and both spheroid detection and classification occur in a single step to streamline the analysis process. Given the fast analysis speed, it’s possible to integrate the Deep-LUMEN analysis in real-time during image acquisition and under live view to guide image selection.

Lumen formation is a gradual process in which a smaller lumen appears first and then gradually enlarges. Although we didn’t quantify the size of the inner lumens of our spheroids over time, we did notice that the degree of confidence automatically provided by the Deep-LUMEN algorithm seems to correlate with the extent of lumen development. Freshly formed lumens with small luminal diameters are often given a confidence of around 85%, while well-developed lumens are assigned a confidence of about 99%. This indicates that even more minute differences in tissue morphology could be discernable. In future studies, rather than limiting 3D model characterization to a two-category classification, additional characteristic parameters, where the gradual change in tissue morphology is transformed into a range of quantifiable values, could be further explored.

## Limitations

The accuracy of our trained model could be further improved with additional training images. However, because much more images can be analyzed with Deep-LUMEN compared to manual analysis, our current level of accuracy didn’t impact the outcome of our experimental analysis, where we can successfully capture the effect of drugs and matrix. Finally, in this study we have only used one type of image cytometer for image acquisition. Image contrast and the filter used could vary between different image cytometers. In future studies, it would be interesting to see if models trained using training images collected from one cytometer can be used for images collected from a different cytometer even when the tissue culture platform is the same. This feature will impact the feasibility of distributing a trained model to different labs with very different imaging infrastructures.

## Conclusions

We have developed a deep learning model, called Deep-LUMEN, that can be used to detect morphological changes in lung epithelial spheroids. We demonstrated that it is possible to develop object detection programs to recognize subtle changes in 3D tissue morphology in response to ECMs and drug treatment. We envision this approach could help accelerate the transition to 3D tissue models for high-throughput drug screening by making the image analysis process less invasive and more informative.

## Materials and Methods

### Cell Culture

A549 cells purchased from Cedarlane labs (Cat# PTA-6231) were cultured in Hams F12K media (Cedarlane labs, Cat# 302004), supplemented with 10% of Fetal Bovine Serum (Wisent Bioproducts, Cat# 098-150), 1% of Penicillin-Streptomycin solution (100X) (Wisent Bioproducts, Cat# 450-201-EL) and 1% of HEPES solution (1M) (Wisent Bioproducts, 330-050-EL). Cells were cultured until 90% confluence in T75 flasks (5% CO_2_, 37 °C). The A549 cells used for all the experiments were between passage 2-5. Before cell seeding, all cells were strained using 40μm cell strainers to remove any cell clumps and get a uniform single-cell suspension.

### Lung spheroid culture

To generate lung spheroids, A549 cells were mixed with growth-factor reduced Matrigel^®^ (Corning, Cat# CACB356231) at a seeding density of 0.1 million cells/ml. 25 µL of this mixture was cast onto one well of a standard 384-well plate (VWR, Cat# 10814-226). For ECM optimization experiments, the cells were also suspended in Fibrin gel. To prepare the fibrin gel, 125 μl of fibrinogen gel aliquot was mixed with 25 μL of thrombin (1.5 U/ml) before gel casting. Both fibrinogen and thrombin stock solutions were prepared as per the supplier’s guidelines (Sigma Aldrich, Cat# F3879-1G, T6884-100UN). Similar to Matrigel^®^ condition, 0.1 million cells/ml were suspended in different concentrations of Fibrin gel (2 mg/ml, 5 mg/ml, and 10 mg/ml), and 25 μL of the final gel mixture was cast into each well. The spheroid culture was maintained for up to 10 days in 5% CO_2_ and 37°C. Culture media was changed every two days.

### Immunofluorescent staining of lung spheroids

Cultured lung spheroids were first washed with 1X PBS to remove residual culture media. The tissue was then fixed with 10% Formalin solution overnight in 4°C. The fixative was removed, and the tissue was washed three times with 1X PBS and blocked overnight in 4°C with 10% Fetal Bovine Serum. The tissue was then stained with F-actin conjugate antibody (Cedarlane Labs, Cat#20553-300) along with DAPI (Sigma Aldrich, Cat#D9542-5MG) and incubated overnight in 4°C. Both antibodies were diluted at 1:200 ratio in PBS with 2% (v/v) FBS. After antibody incubation, the samples were washed in PBS overnight and imaged using an image cytometer.

### TensorFlow Object Detection API Configuration

Image annotations were created using the labeling program. For each image, boxes were drawn around the spheroids and categorized accordingly. The annotations were converted into .xml files containing the bounding box coordinates and the class name for each labeled spheroid. A label map that assigns an integer to each class was also created in a .pbtxt file. Through the use of helper scripts, the annotations were converted into the TFRecord File format. From the TensorFlow detection model zoo, the pre-trained models were downloaded along with their corresponding configuration files. For model 3, 4, and 5: the following changes were made to the configuration file. The number of classes field was set to 2 and the image dimensions field was set to match the images’ dimensions (1224 × 904 pixels). For model 1, and 2, the number of classes was set to 2 and the image dimensions were set to 612 x 452 pixels and 640 x 640 pixels respectively. Finally, for all models, training was set to 200,000 steps. The location of the pre-trained model, the datasets, and the label maps were set.

Model training was conducted using Google Colaboratory. The training was executed on either a Nvidia K80, T4, P4, or P100 GPU, which was randomly assigned with each connection to the Google Colaboratory GPU backend. For model 6, the following data augmentation options were added: random_adjust_brightness, random_adjust_contrast, random_adjust_hue, random_distort color, random_vertical_flip, and random_rotation90. The final lung spheroid training dataset consisted of 3617 images, the validation set consisted of 391 images and the test set contained 197 images.

### Image Acquisition

Lung spheroids were fixed following eight to ten days of culture. The Cytation 5 Cell Imaging Multi-Mode reader (BioTek^®^ Instruments) was used to take z-stack images of the spheroids with a 50 µm step size. Brightfield images were captured under 4X magnification. To capture spheroids from the entire well, images from four different regions of each well were acquired. One well produced 128 total images in .tiff formats. The .tiff images were renamed and simultaneously converted into RGB and .jpg format using a macro script on ImageJ. For fluorescent images and their corresponding brightfield images, the magnification used was 10X.

### Drug administration studies

Cyclosporin A was purchased from Sigma Aldrich (Cat# 30024-25MG). The drug stock solutions were prepared as per the manufacturer’s instructions. The drug was dissolved in Dimethyl sulfoxide (DMSO) (Sigma Aldrich, Cat# D2650-5×5ML) to get a concentration of 25 mg/mL and was then sterile-filtered using a syringe with a filter insert (VWR, Cat# CA28145-501). This solution was diluted 1000 times in culture media to achieve a final DMSO concentration that was lower than 0.1%. Serial dilutions were performed in culture media to get the four concentrations: 10, 1, 0.1, and 0.01 µM. Cyclosporin A was administered for seven days. The drug solutions were refreshed every other day. Z-stack images of 4X magnification were taken every day using Cytation 5 Cell Imaging-multi mode reader.

### Quantification analysis

To evaluate the models based on the number of class instances detected, 197 new test images were chosen and annotated. These served as the ground-truth detections. The detections from each trained model were outputted on these new test images. A detection was considered “correct” if the spheroid was labeled in the ground-truth annotation, and the label matched the ground-truth classification. To measure time to output detections, a human annotator and the model were timed labeling 25 images. The mean Average Precision (mAP) at a 50% intersection of union, using the Pascal VOC performance metrics, was used to measure accuracy. To obtain mAP, a test set was created and each model was analyzed with this test set. To measure the effect of extracellular matrices on spheroid diameter, at least six in-focus spheroids from each condition on day 1, 6, and 10 of culture were measured using ImageJ. To determine the effect of CsA on spheroid diameter, the x-axis bounding box coordinates for all detections above a 50% score confidence were outputted and subtracted from each other. This was done for at least three wells. To assess the effect of CsA on the total number of spheroids and the percentage of lumen spheroids, the width of the bounding boxes were filtered by class and extracted for three independent samples.

### Statistical Analysis

All data are expressed as mean ± standard deviation. Statistical tests conducted were either one-way ANOVA, two-way ANOVA, or one-way ANOVA on ranks followed by the Holm-Sidak or Dunn’s method. For one-way and two-way ANOVA, requirements of normality and equal variance were met. Results were considered significant when a p-value of less than 0.05 was obtained.

## Data Availability

All the trained models and the datasets can be downloaded from https://osf.io/g2a7r/

## Acknowledgments

This work was funded by the National Sciences and Engineering Research Council of Canada (NSERC) Undergraduate Research Award to LA, and Canadian Institute of Health Research (CIHR) Project Grant (PJT-166052) to BZ. The authors are grateful to all the contributors to the Tensorflow Object Detection API and the contributors to BioRender.com which we have used to make the illustrations in this work. Lastly, the authors are grateful to all the essential frontline workers fighting COVID-19.

## Author contribution

L.A. performed the experiments, deep learning analysis, and prepared the manuscript. S.R. contributed to the experiments and prepared the manuscript. D.S.Y.L. contributed to the fluorescent staining of the spheroids. S.V.R and Y.F. contributed to image annotation and quantitative analyses. A.S. and A.L. contributed to image acquisitions. B.Z. envisioned the concept, supervised the work and prepared the manuscript.

## Table of contents

Deep-Learning Uncovered Measurement of Epithelial Networks (Deep-LUMEN) is an open-source object detection algorithm that has been fine-tuned to automatically uncover subtle differences in epithelial spheroid morphology from brightfield images. This algorithm can track changes in the luminal structure of 3D spheroids and distinguish between polarized and non-polarized lung epithelial spheroids. Deep-LUMEN could open new possibilities for assessing drug effects in 3D tissue models or be further expanded to conduct more complex phenotypic screens.

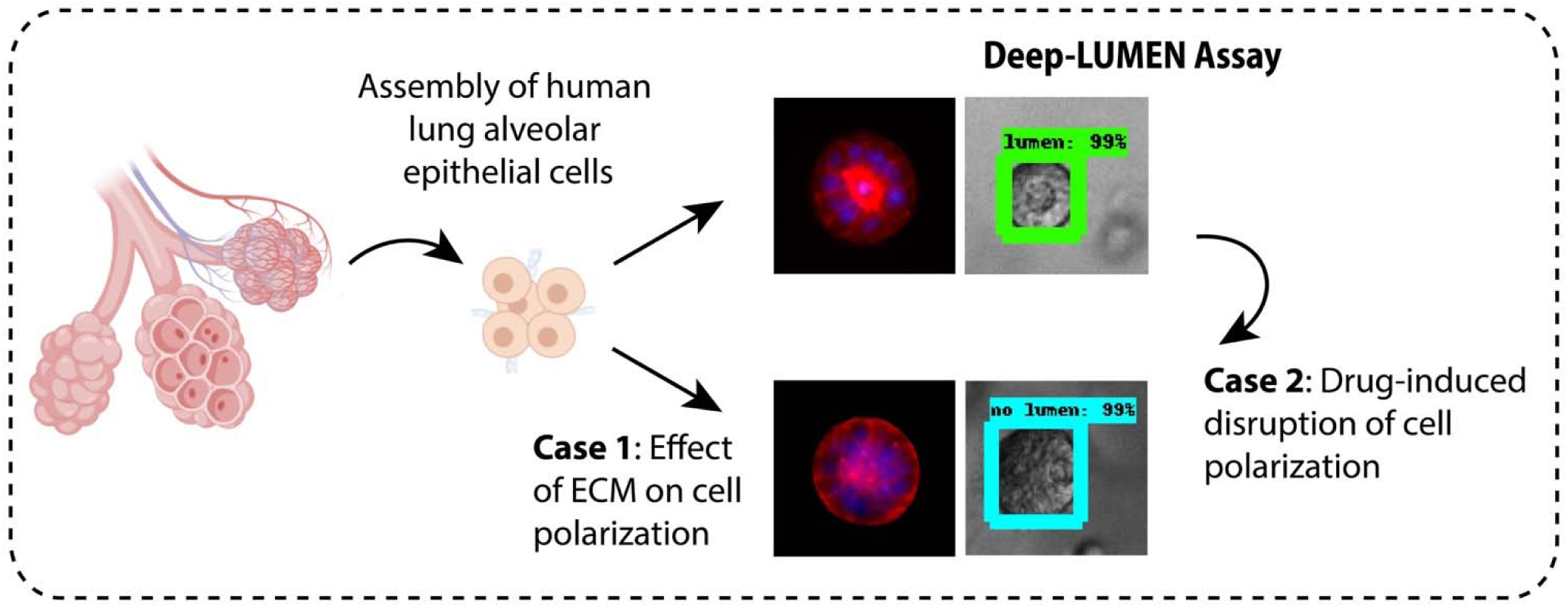

## Supplementary Materials

**Supplementary Figure 1.**
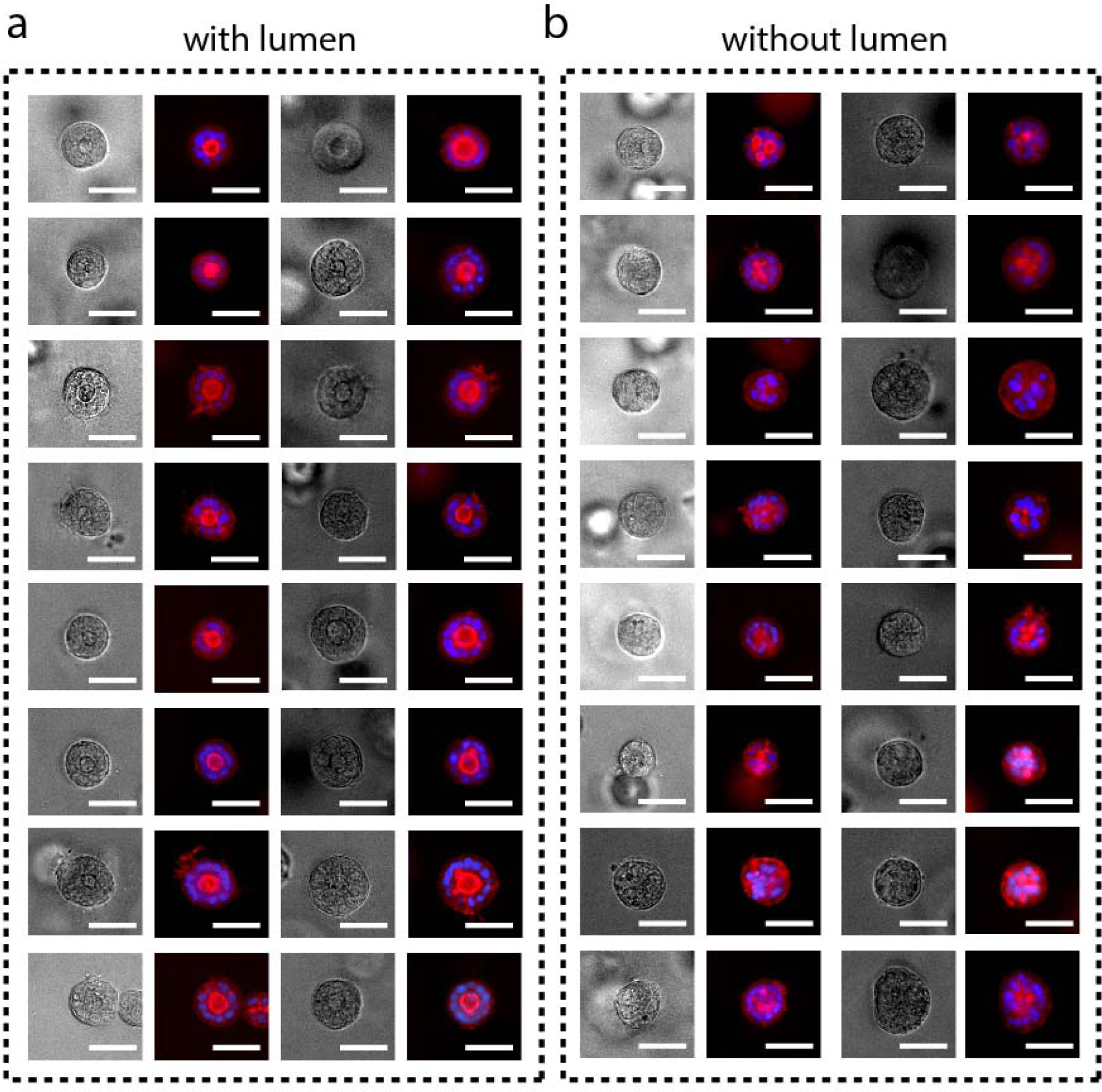
Immunofluorescent staining of polarized and non-polarized lung spheroids. **a-b**, Fluorescent staining of (a) polarized spheroids referred to as “lumen” (representative images n = 6 samples) and (b) non-polarized spheroids referred to as “no lumen”. Spheroids were stained for F-actin (red) and DAPI (blue). Scale bar, 50 μm.

## References

1 Knouse, K. A., Lopez, K. E., Bachofner, M. & Amon, A. Chromosome Segregation Fidelity in Epithelia Requires Tissue Architecture. Cell 175, 200-211.e213 (2018).

2 Bierwolf, J. et al. Primary rat hepatocyte culture on 3D nanofibrous polymer scaffolds for toxicology and pharmaceutical research. Biotechnology and bioengineering 108, 141–150 (2011).

3 Von Der Mark, K., Gauss, V., Von Der Mark, H. & Müller, P. Relationship between cell shape and type of collagen synthesised as chondrocytes lose their cartilage phenotype in culture. Nature 267, 531–532 (1977).

4 Yamada, K. M. & Cukierman, E. Modeling tissue morphogenesis and cancer in 3D. Cell 130, 601–610 (2007).

5 Kota, S. et al. A novel three-dimensional high-throughput screening approach identifies inducers of a mutant KRAS selective lethal phenotype. Oncogene, 1 (2018).

6 Low, J. H. et al. Generation of Human PSC-Derived Kidney Organoids with Patterned Nephron Segments and a De Novo Vascular Network. Cell Stem Cell, doi:https://doi.org/10.1016/j.stem.2019.06.009 (2019).

7 Matano, M. et al. Modeling colorectal cancer using CRISPR-Cas9–mediated engineering of human intestinal organoids. Nature medicine 21, 256 (2015).

8 Qian, X. et al. Brain-region-specific organoids using mini-bioreactors for modeling ZIKV exposure. Cell 165, 1238–1254 (2016).

9 Takebe, T. et al. Vascularized and functional human liver from an iPSC-derived organ bud transplant. Nature 499, 481 (2013).

10 Bayramoglu, N. et al. in 2014 22nd International Conference on Pattern Recognition. 3345-3350 (IEEE).

11 Kecheril Sadanandan, S., Karlsson, J. & Wahlby, C. in Proceedings of the IEEE International Conference on Computer Vision Workshops. 36–41.

12 Kassis, T., Hernandez-Gordillo, V., Langer, R. & Griffith, L. G. orgaQuant: Human intestinal organoid Localization and Quantification Using Deep convolutional neural networks. Scientific reports 9, 1–7 (2019).

13 Soetje, B., Fuellekrug, J., Haffner, D. & Ziegler, W. H. Application and Comparison of Supervised Learning Strategies to Classify Polarity of Epithelial Cell Spheroids in 3D Culture. Frontiers in Genetics 11, 248 (2020).

14 Fathi, A. et al. Speed and accuracy trade-offs for modern convolutional object detectors. (2017).

15 Dai, J., Li, Y., He, K. & Sun, J. in Advances in neural information processing systems. 379–387.

16 Ren, S., He, K., Girshick, R. & Sun, J. in Advances in neural information processing systems. 91–99.

17 Liu, W. et al. in European conference on computer vision. 21–37 (Springer).

18 Lin, T.-Y., Goyal, P., Girshick, R., He, K. & Dollár, P. in Proceedings of the IEEE international conference on computer vision. 2980–2988.

19 O’Brien, L. E. et al. Rac1 orientates epithelial apical polarity through effects on basolateral laminin assembly. Nature cell biology 3, 831–838 (2001).

20 Nussenblatt, R. B. & Palestine, A. G. Cyclosporine: immunology, pharmacology and therapeutic uses. Survey of ophthalmology 31, 159–169 (1986).

21 Wells, G. A. et al. Cyclosporine for treating rheumatoid arthritis. Cochrane Database of Systematic Reviews (1998).

22 Ellis, C. N. et al. Cyclosporine improves psoriasis in a double-blind study. Jama 256, 3110–3116 (1986).

23 Gerets, H. et al. Characterization of primary human hepatocytes, HepG2 cells, and HepaRG cells at the mRNA level and CYP activity in response to inducers and their predictivity for the detection of human hepatotoxins. Cell biology and toxicology 28, 69–87 (2012).

24 Homan, K. A. et al. Bioprinting of 3D convoluted renal proximal tubules on perfusable chips. Scientific reports 6, 34845 (2016).

